# Detecting RyR clusters with CaCLEAN: influence of spatial distribution and structural heterogeneity

**DOI:** 10.1101/549683

**Authors:** David Ladd, Agne Tilunaite, Christian Soeller, H. Llewelyn Roderick, Edmund Crampin, Vijay Rajagopal

**Author notes:** The University of Melbourne, Parkville VIC 3010 Australia.

## Abstract

In cardiomyocytes, coordinated calcium release from intracellular stores through ryanodine receptor (RyR) clusters is key to contraction. Recently, a deconvolution algorithm (CaCLEAN) was proposed to detect the functional response of RyR clusters from confocal imaging of live cells. However, CaCLEAN cluster detection remained unvalidated without ground truth values. We developed a structurally realistic computational model of calcium emanating from RyR clusters in a rat ventricular cardiomyocyte during the first 30 ms of the calcium transient. The effects of RyR cluster density and mitochondria acting as diffusion barriers were examined. Confocal microscopy images were simulated and analysed using CaCLEAN. CaCLEAN detection accuracy was more sensitive to RyR cluster density than the presence of mitochondria. Detection recall and precision varied between 0.69 and 0.82 in densely and sparsely-packed RyR cluster distribution cases, respectively. This sensitivity to cluster packing also affected the distance from the imaging plane clusters were detected.

## Introduction

Each heartbeat is induced by a flood of calcium (Ca2+) from the sarcoplasmic reticulum (SR), into the cytosol. This release of Ca2+ through clusters of ryanodine receptors (RyRs) raises the bulk cytosolic Ca2+ concentration from 0.1 *μ*M to ≈1 *μ*M within 30 ms and results in exposing of cross-bridge binding sites on the actin filaments, facilitating the cellular contractions that compose a heartbeat (Bers, 2002). Confocal (Soeller and Cannell, 2002) and super-resolution (Hou et al., 2015) microscopy imaging of immuno-labelled cardiac tissue preparations has previously revealed the spatial organisation of RyR clusters, but measurements from fixed tissues are limited in their ability to provide insights into function.

Tian et al. (2017) recently proposed an adaption of the CLEAN method from radio astronomy (Högbom, 1974) to detect Ca2+ release sites from fluorescence confocal imaging of live cardiomyocytes. Using this algorithm (CaCLEAN) the authors demonstrated the possibility of detecting the time-dependent functional response of RyR clusters in live cell preparations using widely available experimental methods. Further, if proven in this context, similar methods may extend to other cellular processes that can be approximated as an impulse response of a collection of point sources in a fluorescence microscopy signal.

Several questions remain however regarding the suitability and performance of CaCLEAN. In their original article, Tian et al. (2017) state that based on analysis of Ca2+ transients with varying distance and convolved noise, CaCLEAN is able to resolve local transients with centres 1 *μ*m apart. The authors deem a 1 *μ*m minimum cluster spacing reasonable on the basis that the mean distance between RyR clusters was previously reported as 1.02 *μ*m (Soeller et al., 2007). However, the cited measurement combines clusters within the same z-disk and those spanning separate z-disks. Clusters within a given z-disk are more closely packed, as apparent from the 0.66 *μ*m nearest neighbour distance reported in the same study (Soeller et al., 2007). In the longitudinal orientation typical of experimental preparations of cardiomyocytes (as simulated below in Figure 3A), this results in the presence of inter-cluster spacings below 1 *μ*m in the imaging plane (i.e., along the y-axis of each z-disk in Figure 3A) and in the through-imaging-plane depth.

Further complicating detection is that, compared to the near-vacuum between astronomical objects, the diffusive volume between clusters of RyRs in cardiomyocytes is heterogeneous, consisting primarily of myofibrils and mitochondria. This is relevant to the local distribution of diffusing Ca2+: while homologues of cell membrane Ca2+ transport channels exist on mitochondria and exhibit a modest buffering effect, Ca2+ flux between the cytosol and intra-mitochondrial space is negligible compared to other cytosolic Ca2+ pathways under physiological conditions (Williams et al., 2013). Given this potential barrier-like effect of mitochondria to diffusing Ca2+, we hypothesized that Ca2+ reflecting against mitochondria could impact the detection performance of CaCLEAN.

In order to test the performance of CaCLEAN, we developed a spatially detailed computational model of an eight-sarcomere section of a cardiomyocyte, which we used to simulate reaction-diffusion of Ca2+ emanating from RyR clusters during the rising phase (first 30 ms) of the Ca2+ transient. Fluorescence microscopy data was simulated from this model and analysed with CaCLEAN. This allowed for the establishment of ground truth values for RyR cluster firing locations and assessment of performance in terms of true positives (hits), false negatives (misses), and false positives (algorithmic artefacts).

Our findings indicate that the presence of mitochondria only marginally impacts the performance of CaCLEAN. However, inter-cluster spacing has a significant impact on detection performance and on how far from the imaging plane clusters are most accurately detected. We estimate the accuracy and precision of the algorithm as between 69-82%, depending on the density of cluster locations and their distance from the imaging plane. Our analysis therefore serves as a reference for future applications of this algorithm, providing quantitative analysis of performance using a physics-based modelling framework with known ground truths.

## Results

### Simulating microscopy data allows for assessment of detection performance with known ground truth values

A spatially detailed finite element (FE) computational model of an eight sarcomere section of a cardiomyocyte was constructed to test the performance of CaCLEAN in detecting RyR clusters. The algorithms used to generate the model (Rajagopal et al., 2015) enable the creation of models with different RyR cluster distributions. The influence of mitochondria and the spatial arrangement of RyR clusters on the performance of CaCLEAN were studied using these models. This resulted in four model permutations:

1. CASE 1 (high cluster density, no mitochondria): A case with high cluster density (N=984, 123 clusters per z-disk) based on statistical analysis of nearest neighbour distributions of clusters from immuno-labelled confocal microscopy data (Figure 1B).
2. CASE 2 (low cluster density, no mitochondria): A case with a relatively lower cluster density (N=408 or 51 per z-disk) and an additional constraint of a minimum spacing of 1 *μ*m spacing between cluster centers.
3. CASE 3 (high cluster density, with mitochondria): Mitochondrial regions (Figure 1C) were included within the computational model, acting as diffusion barriers. RyR cluster distributions were defined as in case 1.
4. CASE 4 (low cluster density, with mitochondria): Mitochondrial regions were included; RyR cluster distributions were defined as in case 2.

**Figure 1.**
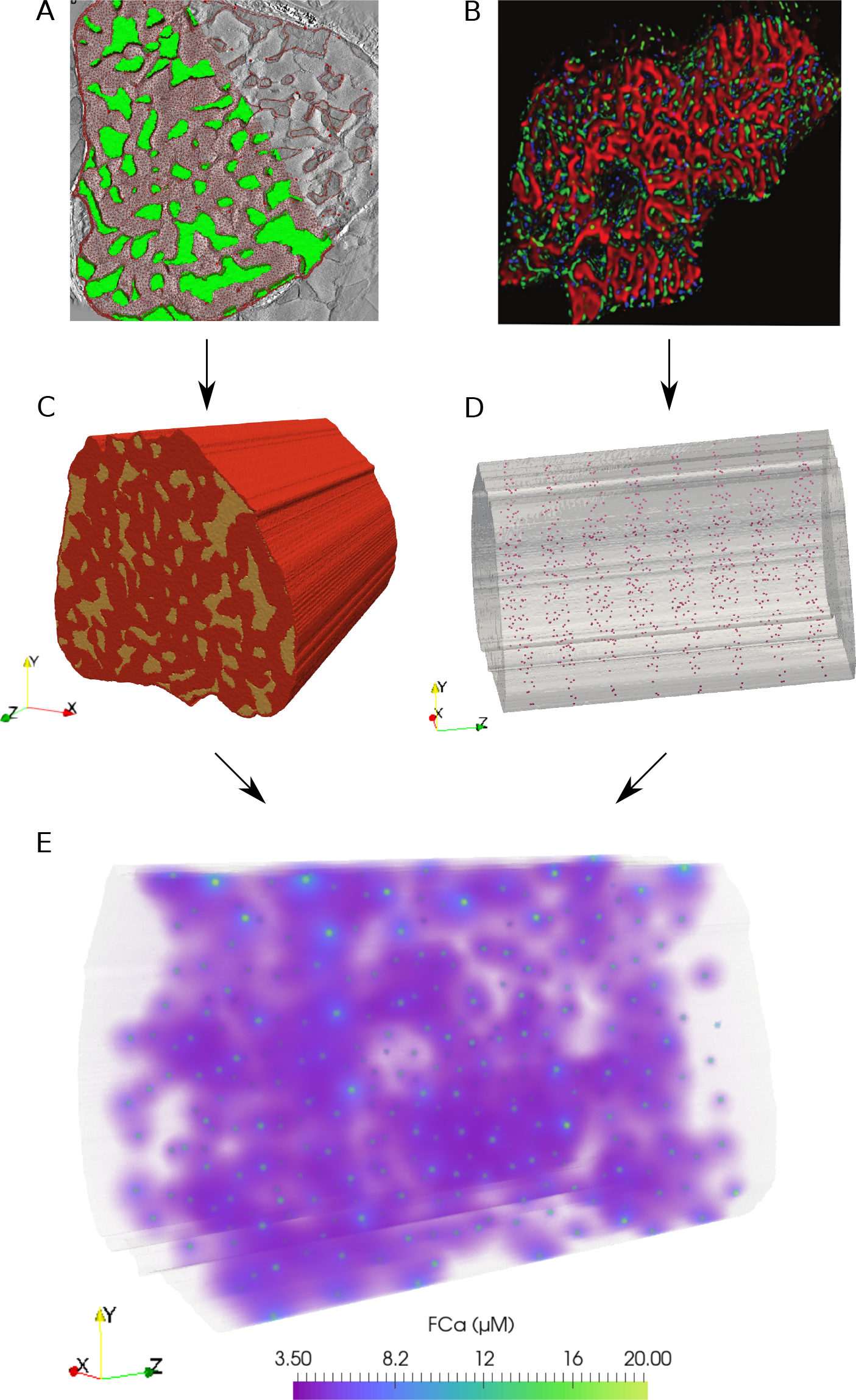
Finite element model of Ca2+ reaction-diffusion in an eight-sarcomere section of a cardiomyocyte. (A) From an electron tomography imaging stack, myofibril and mitochondria (highlighted in green) regions were segmented from a slice at the depth of a z-disk. (C) This geometry was extruded 16 *μ*m (in the direction shown as z here) to create a three-dimensional eight-sarcomere model. Mitochondrial regions shown in yellow; red volume indicates the myofibrillar and cytosolic region. (B) Statistical analysis of immuno-labelled microscopy data (RyR clusters shown in green) was used to determine inter-cluster spacing distributions. (D) RyR cluster locations in the model were defined at mitochondrial and myofibrillar border regions based on statistical spacing distributions. (E) A reaction-diffusion finite element model simulates the release of Ca2+ from the RyR clusters during the rising phase (first 30ms) of the Ca2+ transient. Volume rendering of the fluorescence-bound Ca2+ (FCa) field shown at t = 15ms.

In the three-dimensional FE model, the fluorescence-bound Ca2+ (FCa) field at a given point was determined by the time-dependent reaction-diffusion from the surrounding cluster sources (see Figure 1E). The FCa field was then interpolated onto a regular grid (see Figure 2A) and temporally downsampled to 5 ms intervals. The FCa field was then convolved in three dimensions with a point spread function (PSF, Figure 2B) to simulate optical blurring and light noise (SNR=100) was added. This dataset was then resampled at 215 nm resolution in two dimensions at 22 equidistant slices as indicated in Figure 2C to obtain simulated images that mimic fast 2D confocal images obtained in typical experiments.

**Figure 2.**
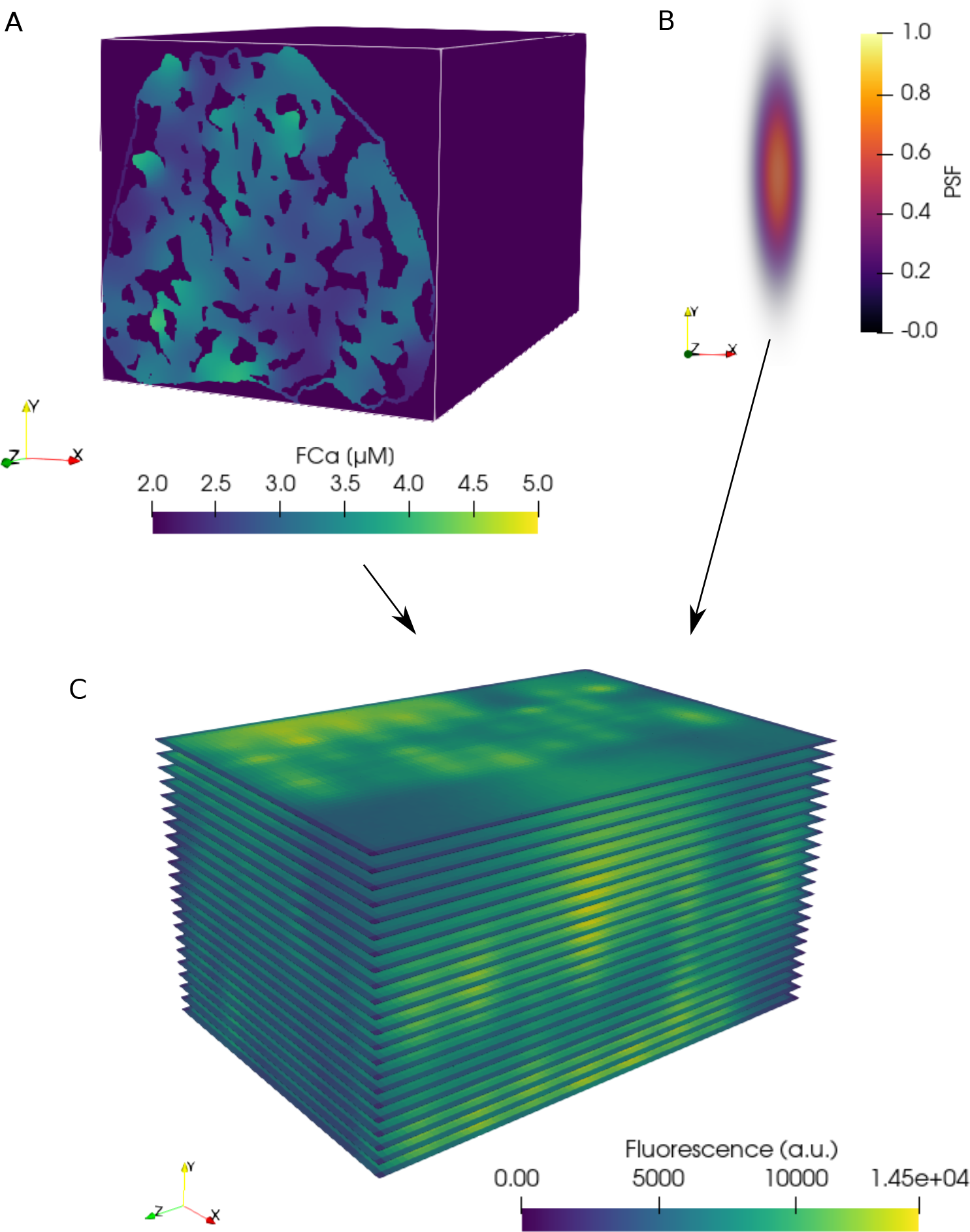
Simulation of confocal fluorescence microscopy images from FE model results. (A) FE FCa field data (see Figure 1E) interpolated onto a regular grid with 53.75 nm resolution in each coordinate direction. Three-dimensional convolution of the interpolated FCa data with a (B) point spread function (PSF) produces blurring typical of confocal fluorescence microscopy data. (C) Blurred data resampled at a pixel resolution of 215 nm within the 22 simulated two-dimensional imaging planes in the volume. A and C are shown at a single time point, t = 15 ms.

The resulting time-dependent, simulated confocal fluorescence microscopy images at each slice (Figure 3A) were then analysed with CaCLEAN to produce maps (Figure 3B), which were then segmented into individual clusters (Figure 3C). To measure algorithm performance we conducted a statistical classification (Figure 3D) using the modeled locations as the actual class (ground truths) and the detected locations as the predicted class to identify true positives (TP), false negatives (FN), and false positives (FP). TP (ground truth) represented the modeled clusters in a given admissible window. TP (detected) represented the TP (ground truth) correctly identified by CaCLEAN. FN identified the TP (ground truth) “missed” by CaCLEAN TP (detected). FP represented the cluster locations identified by CaCLEAN that were not TP (ground truth).

**Figure 3.**
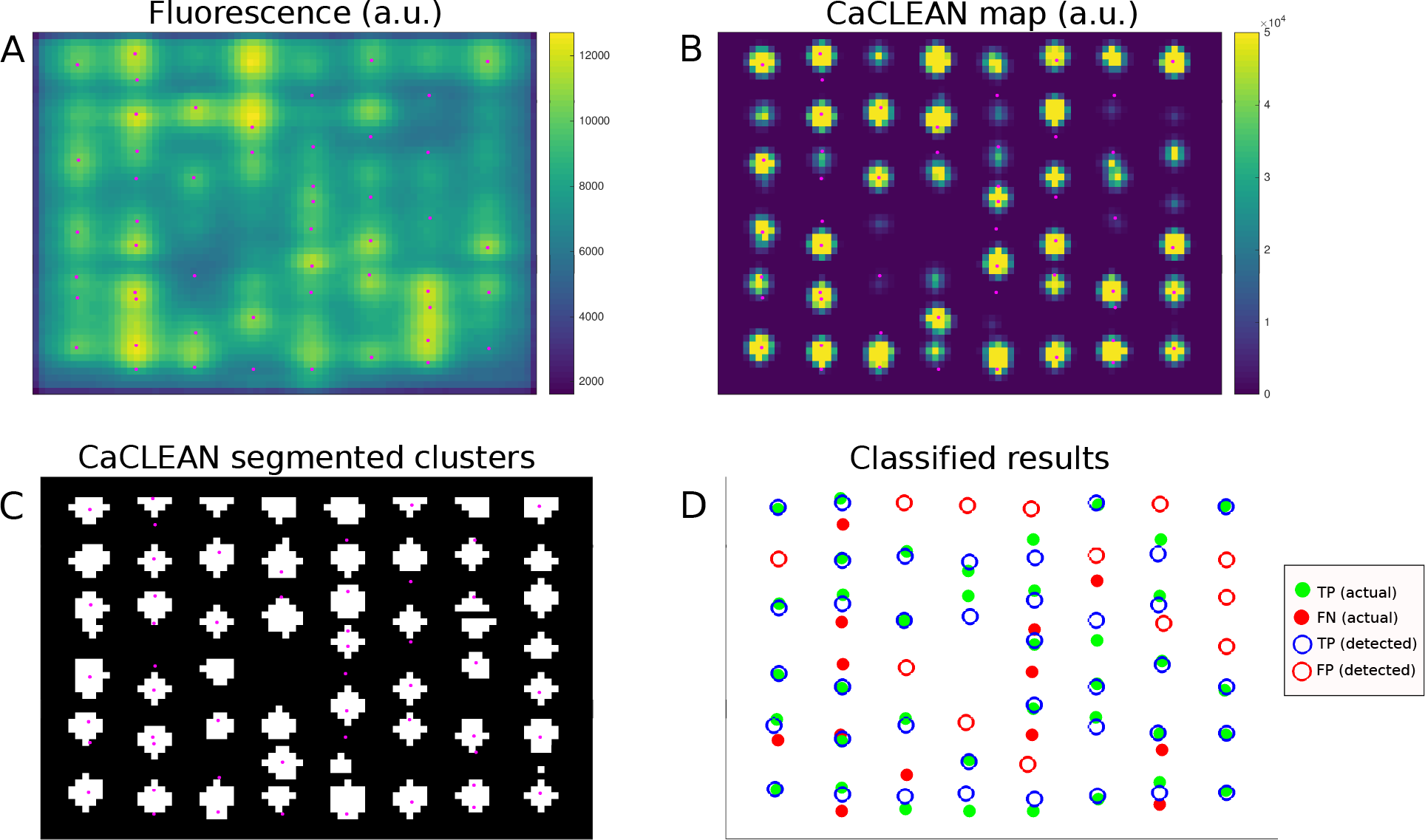
Example of CaCLEAN detection and classification against modeled locations. (A) Simulated confocal fluorescence microscopy image at t=15 ms for the densely packed cluster case (N=123 per z-disk) with mitochondria. This represents one slice from the middle of the stack shown in Figure 2C. Modeled RyR cluster center locations within 280 nm of the imaging plane shown in magenta (also in B and C). (B) CaCLEAN map of the fluorescence signal. (C) Segmented clusters detected by CaCLEAN. (D) Statistical classification of detected cluster locations versus actual (ground truth) modeled locations. In this case recall = 0.68 and precision = 0.68, representing an ‘intersection point’ admissible window, as described further below.

### Cluster distance from the imaging plane reveals the trade-off between recall and precision

For each simulated imaging plane, modeled RyR clusters were considered TP (ground truth) if their centers were within a distance threshold from the imaging plane we referred to as the ‘admissible window’ (see Figure 4). As illustrated in Figure 4A, the number of TP (ground truth, magenta) increased linearly with the admissible window as the window incorporated more of the modeled cluster locations. The number of TP (detected, green) events approached a limit as the collapsed two-dimensional imaging space became saturated with available TP (ground truth) locations and the signal diffusing from far-field clusters did not reach the image space.

**Figure 4.**
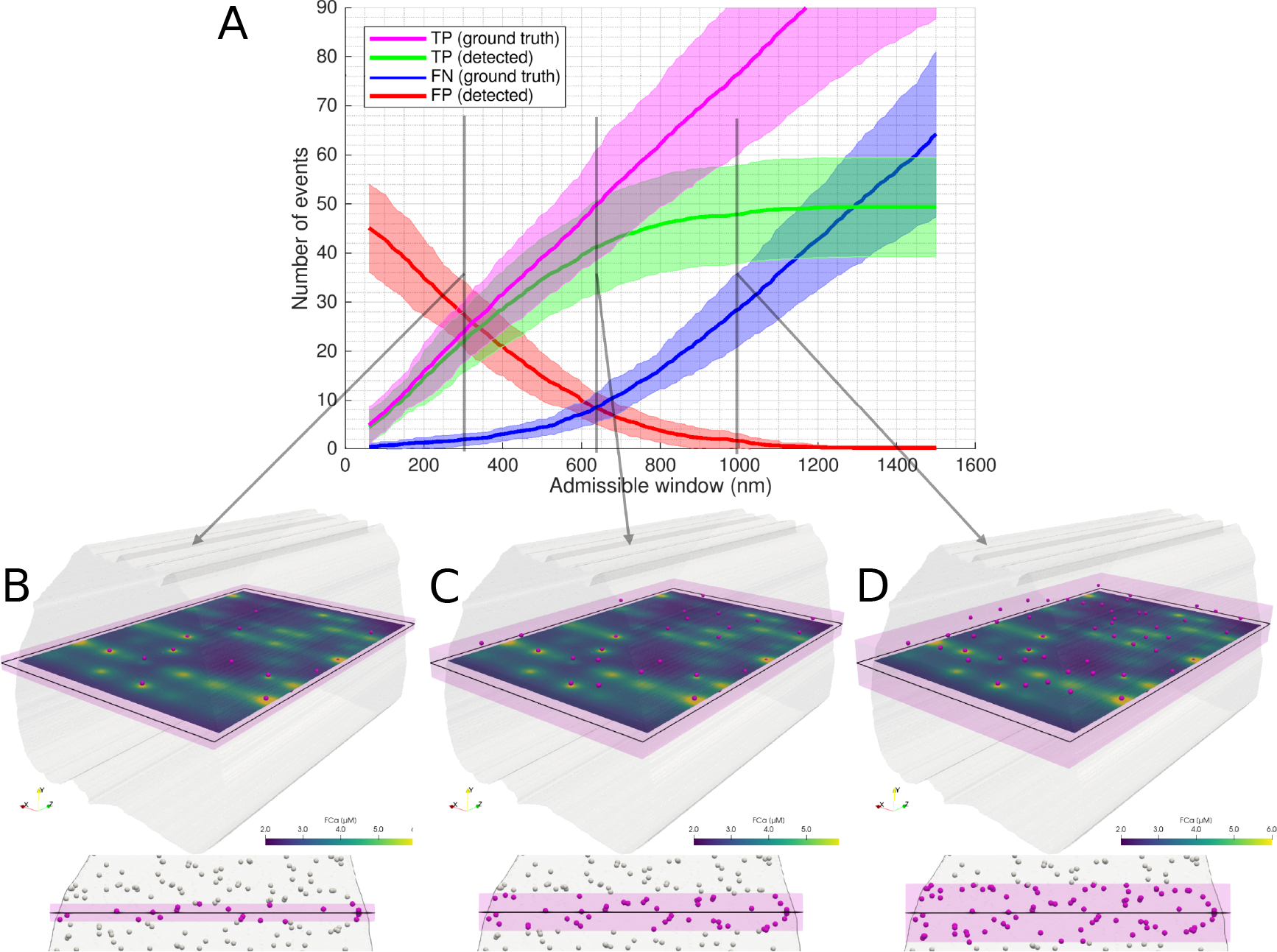
Illustration of the admissible window parameter. The admissible window defines the distance tolerance from the simulated imaging plane RyR cluster centers may be to be considered ground true positives. (A) Statistical classifier CaCLEAN results as a function of the admissible window for low cluster density, no mitochondrial barriers (case 2). 22 images were simulated with equidistant spacing through the model volume. Solid lines represent mean values and shaded regions indicate one standard deviation of values. In (B), (C), and (D) an example image slice is shown, with clusters considered ground TPs at admissible windows of (B) 300 nm, (C) 640 nm, and (D) 1000 nm in magenta. An oblique view is shown above, along with FCa model results at t = 15ms. Below, an axial view perpendicular to the imaging plane is shown (looking through the modeled volume, with clusters outside the admissible window shown in white). The admissible window is indicated in pink shading.

Clusters located very near the z-depth^1^ of the imaging plane were the most likely to affect the signal and be detected by the algorithm, as indicated by the consistently low FN (blue) at low admissible window. However, with a very narrow admissible window, FP (red) were prevalent since clusters located just outside of this arbitrary tolerance still diffused into the imaged space and were detected by CaCLEAN. Conversely, with a widening definition of the admissible window, FP dropped and FN increased due to the asymptotic behaviour of TP (detected).

Recall and precision were evaluated for the four model permutations (see Figure 5). Recall, as the ratio of TP (detected) to TP (ground truth), provided fractional measure of correctly identified clusters. Precision, as the ratio of the number of TP (detected) to all detected sites (TP and FP), provided the fraction of detected clusters that were not FP. Higher values indicate better performance in both measures. Precision appeared to asymptotically approach 1 in all cases except the densely-packed with mitochondrial barriers case (case 3), where additional false positive events resulted in a slight drop to 0.95.

**Figure 5.**
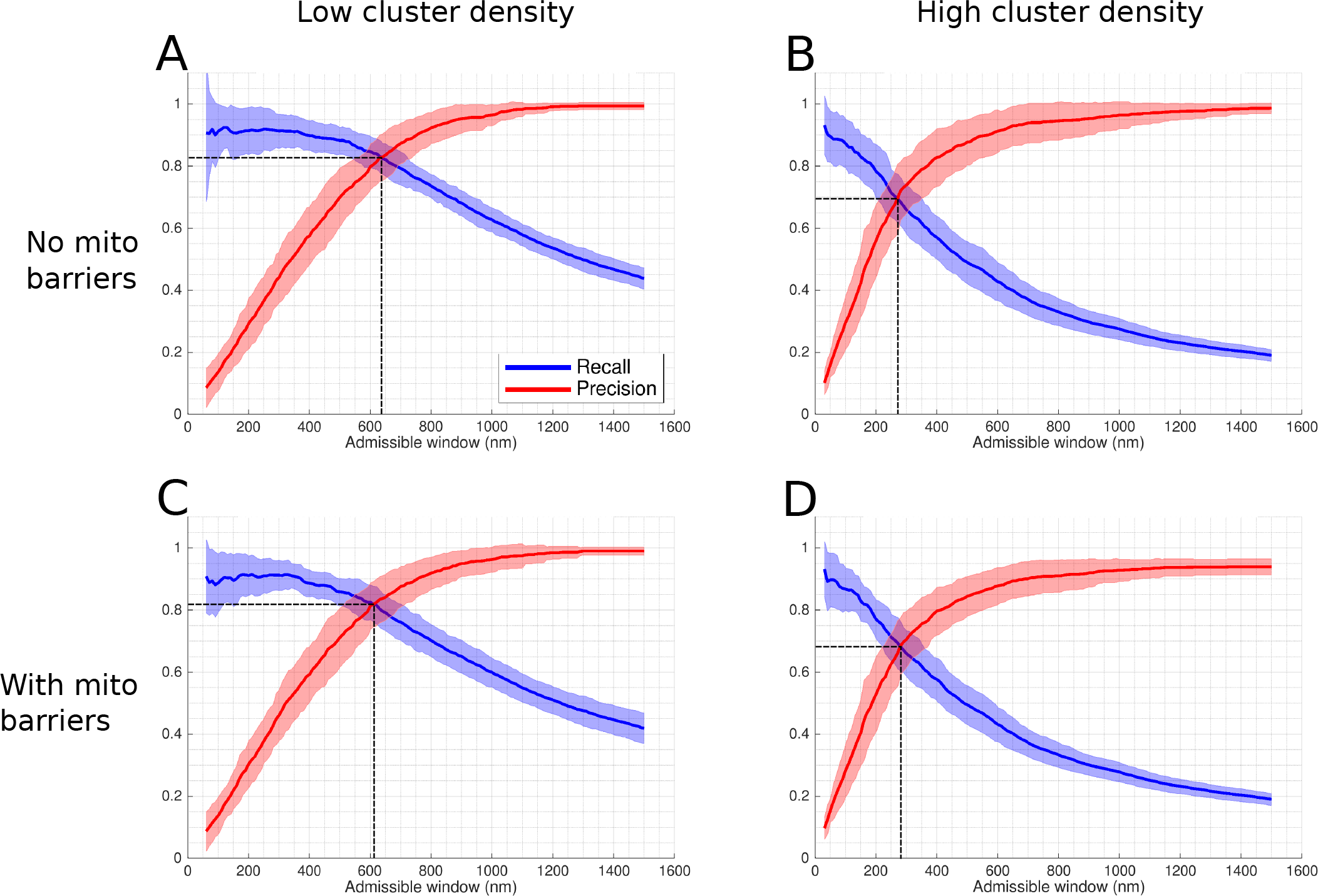
Recall and precision of CaCLEAN applied to simulated microscopy data. Recall and precision evaluated based on classification results. Shaded regions indicate one standard deviation of recall and precision values. (A) A minimum spacing of 1 *μ*m is enforced between modeled clusters, no mitochondria present in the volume (resulting in a continuous domain across the mitochondrial regions). (B) Cluster spacing is determined by statistical analysis of RyR cluster distributions from immuno-labelled super-resolution microscopy data, no mitochondria present in the volume. (C) A minimum cluster spacing of 1 *μ*m is enforced between modeled clusters, mitochondrial regions are subtracted from the volume, acting as barriers to diffusion. (D) Cluster spacing is determined by statistical analysis of RyR cluster distributions from immuno-labelled super-resolution microscopy data, mitochondrial regions are subtracted from the volume, acting as barriers to diffusion. Dashed black lines used to indicate values at ‘intersection points’, where recall = precision.

### Inter-cluster spacing has a greater impact on CaCLEAN performance than mitochondrial diffusion barriers

The intersection of the recall and precision curves identifies the admissible window at which the number of false positives is equivalent to the number of false negatives (and therefore the ‘correct’ number of clusters is still predicted). While there may be applications where optimising for recall or precision may be more appropriate, this intersection point was chosen a useful indicator of general performance. The intersection point values are indicated by the dashed lines in Figure 5 and Figure 6. Without mitochondria, in the low cluster density case recall = 0.83 ± 0.049 and precision = 0.83 ± 0.075 at an admissible window of 640 nm; in the high cluster density case recall = 0.70 ± 0.079 and precision = 0.70 ± 0.112 at an admissible window of 270 nm.

**Figure 6.**
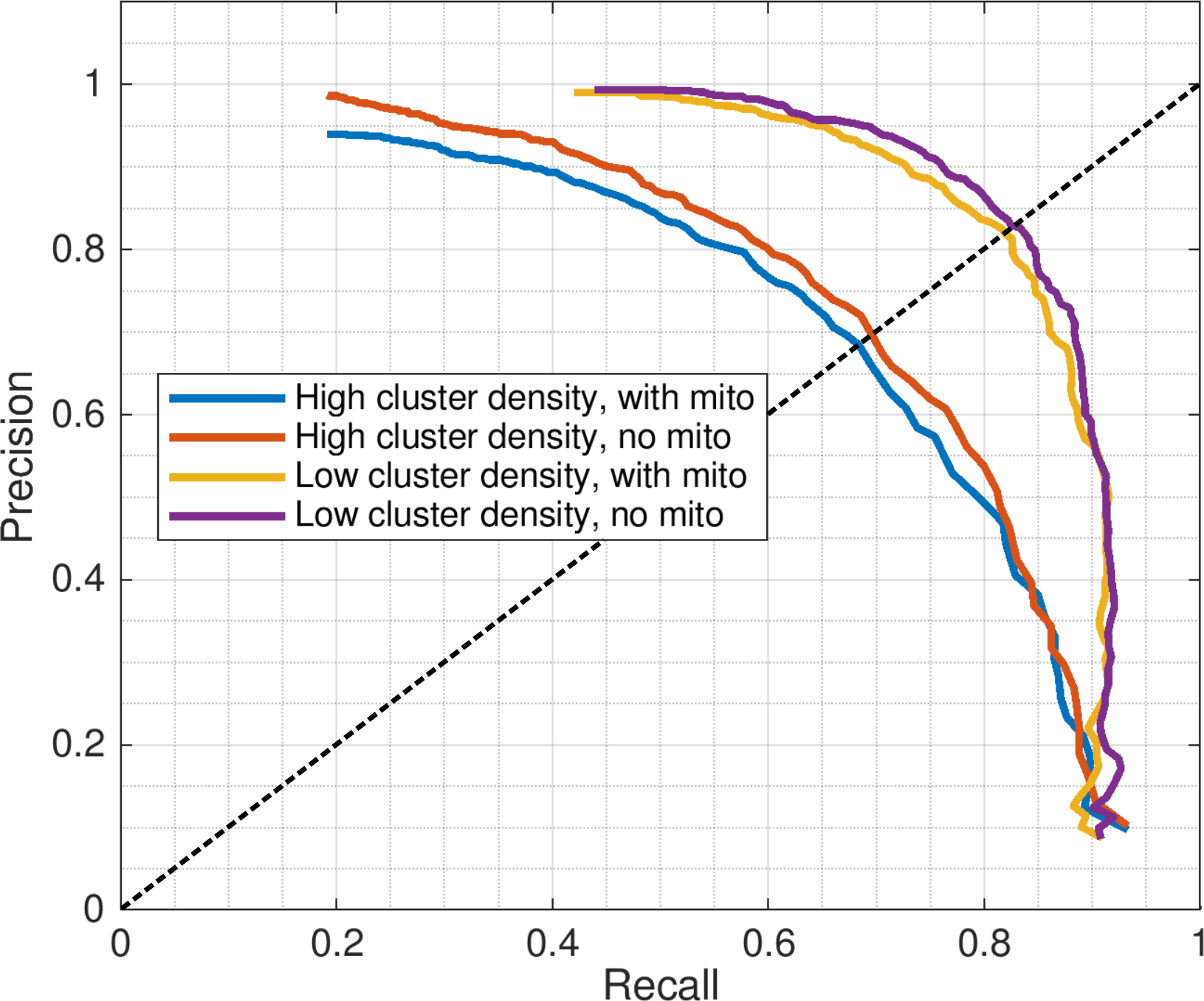
Precision-recall curve for CaCLEAN in four models. Recall and precision mean values from results shown in Figure 5 plotted as a precision-recall curve. Higher values indicate better performance in each axis. Dashed line indicates intersection point, where recall = precision.

Inter-cluster spacing was found to have a greater impact on the performance of CaCLEAN than the presence of mitochondria. With mitochondria acting as barriers to diffusion, in the low cluster density case recall = 0.82 ± 0.059 and precision = 0.82 ± 0.073 at an admissible window of 610 nm; in the high cluster density case recall = 0.69 ± 0.079 and precision = 0.69 ± 0.091 at an admissible window of 280 nm. Mean recall and precision values for each model permutation (solid lines in Figure 5) were further analysed to produce precision-recall curves (see Figure 6).

Finally, we evaluated the fraction of clusters detected by CaCLEAN as a function of z-distance from the focal imaging plane to identify how detection of clusters decayed with distance from the imaging plane (see Figure 7). This analysis again highlighted the effect of inter-cluster spacing: when using Ca-clean to detect sites in models with a high density of release sites the effective z-response of detection falls much more steeply than when a low density of true release sites was simulated.

**Figure 7.**
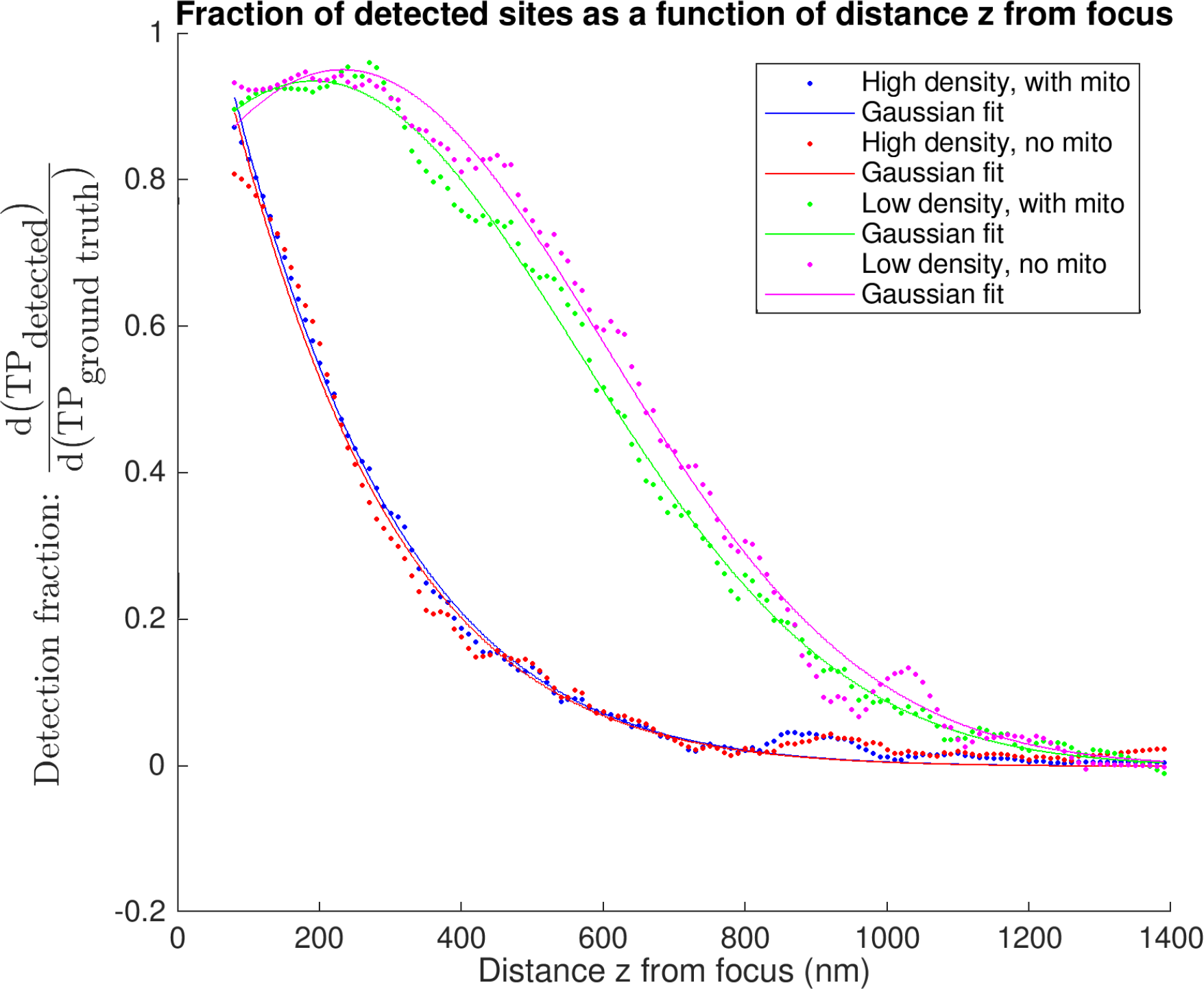
“Differential recall”: axial dependence of detection of true positive sites. Each curve shows the fraction of detected sites as a function of the distance z from the nominal focal plane. The data at focal distance z is shown as a fraction of all sites within 5 nm of that z depth i.e. within a 10 nm band centered about the distance z above and below the focal plane. A value of 1 is equivalent to the detection of all sites at a given z-depth. The various curves are calculated for different models, having a high or low density of sites and either using homogenous diffusion (“no mito”) or obstacles to diffusion wherever mitochondria are (“with mito”). Data points shown are the result of applying a smoothing filter (see Figure Supplement 1 for the low density, no mito case shown in magenta). Solid lines indicate single-term Gaussian fits to the filtered data.

## Discussion

We developed a computational model of the complex environment of Ca2+ diffusing into the intracellular space of a cardiomyocyte. Processing the reaction-diffusion model results to simulate confocal fluorescence microscopy data allowed for quantitative assessment of the performance of detection of Ca2+ release sites with CaCLEAN against known ground truth values in the context of realistic cellular physics. Statistical classification identified true positives, false positives, and false negatives^2^; enabling analysis in terms of recall (sensitivity, hit rate, or true positive rate) and precision (positive predictive value).

A key variable in the performance analysis was the definition of which modeled clusters were considered ground truths in the statistical classification at each imaging plane. We introduced the ‘admissible window’ variable for admitting clusters as ground truth values based on the through-imaging-plane distance of cluster centers from the imaging plane, as illustrated in Figure 4. Parameterising algorithmic performance in terms of the admissible window provided a more complete picture of which clusters CaCLEAN was likely to detect and highlighted the inherent trade-off between precision and recall.

The admissible window is also of practical interest to those applying CaCLEAN to their data. This parameter can be used as an indicator of the maximum relevant depth of clusters detected by the algorithm in the image. As evident in Figure 5, interpretation of this depth is strongly dependent on a user’s performance requirements and the density of clusters in the sample. A user seeking to determine maximum relevant depth should first determine whether they are more willing to sacrifice precision or recall, deciding whether misses or false positives are of greater concern in their application. We suggest the ‘intersection point’ as a general default performance measure: this identifies the depth where precision is equal to recall and the number of false positives is equal to the number of false negatives (thus still detecting approximately the correct number of clusters). For example, if a user is confident that clusters in a sample are likely at least 1 *μ*m apart and is satisfied with correctly identifying at least 80% of the clusters that exist, the maximal relevant depth would be ≈610 nm based on Figure 5C. From the recall curve, the same user may also be interested to find the algorithm will likely correctly identify 90% of all clusters within 350 nm of the imaging plane. The recall curves in Figure 5 therefore also communicate how the detection performance for an active cluster population drops as further away clusters are considered ground truths.

The shape of the point spread function (PSF, see Figure 2B) weights the interpolation and blurring of the three-dimensional data into a two-dimensional image at each time step. Choice of PSF impacts the fidelity of the data interpolation into the image and, subsequently, CaCLEAN’s performance on the interpolated data. However, it should be noted that the optical depth of the PSF does not directly influence how far from the image clusters may be located or detected. CaCLEAN detection operates on the fluorescence signal emitted by Ca2+ released from RyR clusters, rather than directly capturing these sites in the optical depth of the microscope. Figure Supplement 1 shows the recall and precision values using PSFs with half and double the full width at half maximum dimensions of our baseline settings. A tighter PSF produces less blurring and slightly better performance in both precision and recall, while a broad PSF produces excessive blurring and a drop in performance but general trends remain similar.

In addition to distance from the imaging plane, further factors complicate whether the signal from a given cluster will reach the imaging plane and whether it is detectable by CaCLEAN in both our model and actual experimental data. Individual cluster firing time and strength variability biases detection of early and stronger events. CaCLEAN produces some false positives as algorithmic artefacts. Proximity to other clusters can cause signals to merge or cover each other-especially with increased cluster density. For instance, two separate clusters the same z-distance above and below the imaging plane but at the same x-y location in the imaging plane can only register as a single site in the imaged space. This identifies an inherent drawback resulting from using two-dimensional images as a basis for describing a three-dimensional system.

We investigated the impact of cluster spacing on CaCLEAN detection performance by analysing two cluster distribution types. In the ‘high cluster density’ case, cluster locations at each z-disk were defined based on statistical analysis of cluster distributions from immuno-labelled confocal images and applied to admissible locations bordering mitochondria and myofibrils segmented from electron tomography data (Rajagopal et al., 2015). This simulated a case where all RyR clusters detectable by immuno-labelling^3^ fired during the 30 ms period modeled. This was considered a reasonable high bound on cluster distribution density since it is unlikely that all clusters identifiable by immuno-labelling would fire for a single excitation cycle under normal physiological conditions. Tian et al. (2017) used CaCLEAN to estimate cluster re-fire rates of ≈ 62.8% in mouse atrial myocytes, with only ≈ 10% of clusters always firing. However, the authors also showed that beta-adrenergic stimulation can increase recruitment of firing clusters, reporting increases in detected cluster density of ≈ 30% in rat ventricular myocytes^4^. In our ‘low cluster density’ case, distributions of clusters were similarly generated based on realistic locations of RyR clusters but with an additional constraint of a minimum spacing of 1 *μ*m between all clusters within a given z-disk. The number of clusters in this case was also reduced to comply with this constraint, with the total number of clusters 41% of those of the high density case. This was chosen as representative of a lower bound on cluster recruitment (as might occur with high pacing frequency) but also revealed the impact of a 1 *μ*m minimum cluster spacing requirement on detection.

RyR cluster distribution spacing varies across species, with nearest-neighbour spacings reported as 0.66 ± 0.06 *μ*m in rat and 0.78 ± 0.07 *μ*m in human (Soeller et al., 2007). Clusters located in the periphery of mouse myocytes are more irregularly spaced than those located in the cell interior (Hiess et al., 2018). These distributions also alter during development, with RyR clusters in rabbits changing from majority peripheral clusters with ≈0.7 *μ*m spacing in neonates to majority interior clusters with ≈2 *μ*m regular spacings between z-disks (Dan et al., 2007). In addition to spatial locations of RyR protein clusters, another consideration is functional response of clusters; an area where CaCLEAN shows unique promise. Tian et al. (2017) used the algorithm to explore how firing reliability decreases with increased stimulation frequency and how firing reliability increases under beta-adrenergic stimulation.

In our analysis, cluster spacing had a significant impact on both the performance of CaCLEAN and the admissible window size at which the algorithm performed best. At the intersection point of the ‘with mitochondria’ cases, detection recall and precision were 0.82 at 610 nm in the low cluster density case versus 0.69 at 280 nm in the high cluster density case. In contrast, for the low cluster density results at the high cluster density intersection point admissible window of 280 nm, precision = 0.43 indicating more false positives than true positives.

Analysis of performance in terms of the admissible window provides a basis for assessing “cumulative recall”, i.e., the detection of all clusters within a given tolerance of the focal plane. We also evaluate “differential recall” in Figure 7, i.e., the detection fraction of clusters at distance z from the imaging plane. This communicates how the detection fraction of individual clusters drops with distance from the focal plane and again emphasizes the importance of cluster density: detection falls much more steeply in the high density cases. In other words, depending on how many sites are active the z-response of CaCLEAN changes. In experiments this means that in cases where all sites release, e.g. after stimulation with with a drug such as isoprenaline, the sites reported by CaCLEAN are on average from regions closer to the focus than when recording in conditions of partial block where fewer sites are available.

For useful detection performance, it is therefore important to consider the likely density of the events being detected in order to determine how far from the imaging plane such events are likely located. Those interested in using CaCLEAN to reconstruct three-dimensional maps of clusters should also be aware of how far from the imaging plane detected clusters are likely to be when choosing a slicing depth for reconstruction.

The presence of heterogeneous diffusion (in the form of mitochondrial obstacles) in the investigated models was found to have a marginal but consistent negative impact on both recall and precision in CaCLEAN. This was most pronounced in the high cluster density case, as evident from the precision-recall analysis (Figure 6). We hypothesize that structural heterogeneity due to the presence of subcellular structures (e.g., mitochondria, nucleus, transverse tubules, z-disks, etc.) is unlikely to have a major impact on detection of point sources using CaCLEAN or related algorithms under the imaging resolution conditions simulated here.

The results presented in this work (particularly Figure 5, Figure 6, and Figure 7) are provided as a reference tool for potential users of CaCLEAN. The model and the simulated imaging data produced by it could also serve as a testing and validation platform for further improving CaCLEAN or other approaches seeking to identify RyR clusters by serving as a training set for improved detection or segmentation. Finally, this study highlights how computational methods may be used to establish ground truth values that may not otherwise be experimentally available.

## Materials and methods

### Finite element model of reaction-diffusion in a half-sarcomere

The reaction-diffusion finite element (FE) model here builds on a previous FE model of a half-sarcomere (Rajagopal et al., 2015). From electron tomography (ET) images of adult rat ventricular myocytes, a three-dimensional axial region with a thickness of approximately 0.875 *μ*m was segmented. This region represented approximately half of a single sarcomere, with the z-disk a plane through the center of the thickness of the domain. The region was approximately 11 *μ*m in diameter, varying with the segmented surface.

From this half-sarcomere image stack, the central slice representing the level of the z-disk was extracted and regions representing myofibrils and mitochondria were manually segmented. This two-dimensional slice was then extruded 16 *μ*m to create a three dimensional volume.

Two configurations were considered to assess the impact of mitochondria: one in which the interior of the modeled domain is a homogeneous material continuum and one in which the regions representing mitochondria were subtracted from the myofibrillar and cytoplasmic domain. The latter case was based on the assumption that the calcium buffering activity of mitochondria is negligible.

Ryanodine receptor (RyR) locations were determined algorithmically, using a spatial statistics method based on nearest neighbour distances of experimentally-derived RyR locations (Rajagopal et al., 2015). These distributions were determined from confocal images of left ventricular cardiomyocytes of a healthy adult male Wistar rat using multiple passes of a band-pass filter detector, following the technique described by Soeller and Cannell (2002). The method also limits the admissible locations of RyR clusters to segmented regions bordering myofibrils, mitochondria, and the sarcolemma (i.e., RyR clusters were not placed inside these organelles).

To evaluate the influence of spacing on the RyR cluster detection performance of CaCLEAN, two sets of constraints on the RyR distributions were considered: (1) RyR clusters were assigned following Rajagopal et al. (2015), with locations based directly on statistical analysis of experimental data and with the total number of RyR clusters N=123 per z-disk (984 total); (2) a minimal distance constraint of 1 *μ*m between RyR cluster centers was enforced and the total number of clusters was reduced to N=51 per z-disk (408 total) to allow for this constraint. Eight unique cluster distributions were generated for each case and were spaced 2 *μ*m apart along the extruded axis model volume to represent z-disk RyR populations. Linear tetrahedral meshes were constructed on these domains, consisting of: (1) 1436943 nodes, 8222684 elements; and (2) 1318942 nodes, 7504655 elements respectively. These meshes included increased refinement in the regions containing and surrounding the modeled RyR locations.

Spherical regions 100 nm in radius were defined around nodes nearest to the determined location for each RyR cluster. Nodes lying within this sphere were prescribed density amplitudes exponentially decreasing as a function of the square of their radial position from the central node. RyR cluster release times were sampled from an exponential distribution with a characteristic decay constant of 6.7 ms, following the findings of Wang et al. (2001).

Further details on the reaction diffusion equations for the buffers modeled and the ordinary differential equation model describing release of Ca2+ from RyR clusters have been previously reported (Rajagopal et al., 2015).

### Simulation of confocal fluorescence signals from FE model results

To simulate confocal fluorescence results, the irregularly distributed node-based fluorescence-bound Ca2+ field from the FE model was first interpolated onto a regular grid at a resolution of 53.75 nm in each direction. The data was temporally sampled at 5 ms intervals over the simulated time period of 30 ms, producing simulated imaging data for 7 timesteps. Nodal positions within mitochondria in the ‘with mitochondria’ permutations were ascribed the initial and background value FCa = 2.08. Discrete natural neighbor (Sibson) interpolation (Park et al., 2006) was chosen for this task on the basis that it can be used to generate regularly-spaced three-dimensional data, scales well for large datasets, and does not require additional parameterisation. The implementation used was version 1.7 of the naturalneighbor python package, available from the Python Package Index under the MIT license.

A point spread function (PSF) was generated as a normalised function of a multivariate Gaussian distribution applied in three dimensions (see Figure 2B). These distributions had a full width at half maximum (FWHM) of 410 nm in x and y and 1800 nm in z (where the z-axis represents the through-imaging-plane direction, shown as ‘y’ in Figure 2B.). These values were based on reported estimates of the dimensions of a PSF from a Visitech confocal microscope (Plumb et al., 2015). The resolution of the PSF image was chosen to be the same as the interpolated model data (53.75 nm in each direction).

The interpolated model imaging data was then convolved with the PSF in three dimensions using the SciPy convolve algorithm from the signal processing module. The resulting grid was then downsampled to a pixel resolution of 215 nm, following the resolution of the original CaCLEAN paper (Tian et al., 2017). Light noise (SNR = 100) was also added to the image data. Representative slices were sampled along the y-axis of the FE model, producing 22 two-dimensional simulated microscopy images for each of the four reported model permutations.

### Application of CaCLEAN to simulated fluorescence data

The CaCLEAN algorithm was obtained from the author’s GitHub repository: https://github.com/qhtian/CaCLEAN. The Matlab-based scripts were run using Matlab version R2017b. The function CICRcleanSimp was used to generate the CaCLEAN release map and the function CRUProps was used to segment the release map into individual calcium release units (CRUs).

### Classifying and quantifying performance of CaCLEAN

A statistical classification approach was used to assess the performance of RyR cluster detection using CaCLEAN. Modeled cluster centers within the admissible window were considered the actual / ground truth class: TP (ground truth). CaCLEAN detection results were considered the predicted class. Detected RyR cluster sites were defined as determined by the CaCLEAN CRUProps function, which segments cluster regions using Matlab’s built-in watershed algorithm and identifies centroids of segmented regions.

For each modeled cluster location, a TP (detected) classification was assigned if a TP (ground truth) cluster center lied within an available segmented CaCLEAN cluster region (or within a 1 pixel tolerance). When more than one TP (ground truth) fell within a CaCLEAN-detected cluster region, the detected cluster with the nearest centroid to the TP (ground truth) location was marked TP (detected). After classification as TP (detected), the associated CaCLEAN-detected site would be removed from the list of available matches. After iterating through the TP (ground truth) clusters, remaining TP (ground truth) unmatched with detected clusters were classified as false negatives (FN). Remaining CaCLEAN-detected sites unmatched with TP (ground truth) were classified as false positives (FP).

Two statistical measures were considered: recall and precision. Recall (also known as sensitivity, hit rate, or true positive rate) was defined such that

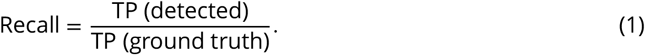

Recall therefore gives the fraction of actual modeled clusters within an admissible window that were correctly detected by CaCLEAN. Precision (also known as positive predictive value) was defined such that

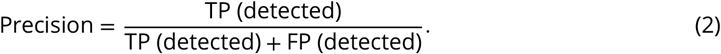

Precision therefore identifies the fraction of the detected clusters within an admissible window that were correct (not false positives). In both cases, higher values indicate better performance.

The above definition of recall may be considered “cumulative recall”, identifying the detection fraction of all clusters within a given admissible window. To determine the detection fraction of clusters at a given z distance from the simulated imaging plane, we also defined an alternative “differential recall” such that

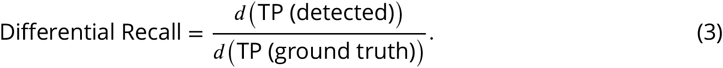

This measured the fraction of modeled clusters detected by CaCLEAN in 10 nm spaced bands above and below the imaging plane. Only bands with at least one TP (ground truth) were considered. Mean values for this detection fraction were acquired over the 22 simulated imaging planes. A Savitzky-Golay filter (polynomial order 3, frame length 21) was then applied to smooth the results as shown in Figure Supplement 1. Single-term Gaussian fits were also applied to identify trends in the resulting curves, as shown in Figure 7.

## Acknowledgments

This research was supported in part by the Australian Government through the Australian Research Council’s Discovery Projects funding scheme (project DP170101358), and in part by the Australian Research Council Centre of Excellence in Convergent Bio-Nano Science and Technology (project number CE140100036). HLR wishes to acknowledge financial support from the Research Foundation Flanders (FWO) (Project Grant G08861N and Odysseus programme Grant 90663). CS acknowledges financial support by the Engineering and Physical Sciences Research Council of the United Kingdom (Grant EP/N008235/1) and Biotechnology and Biological Sciences Research Council Grants BB/P026508/1 and BB/R022127/1.

---

**Figure 4–Figure supplement 1.**
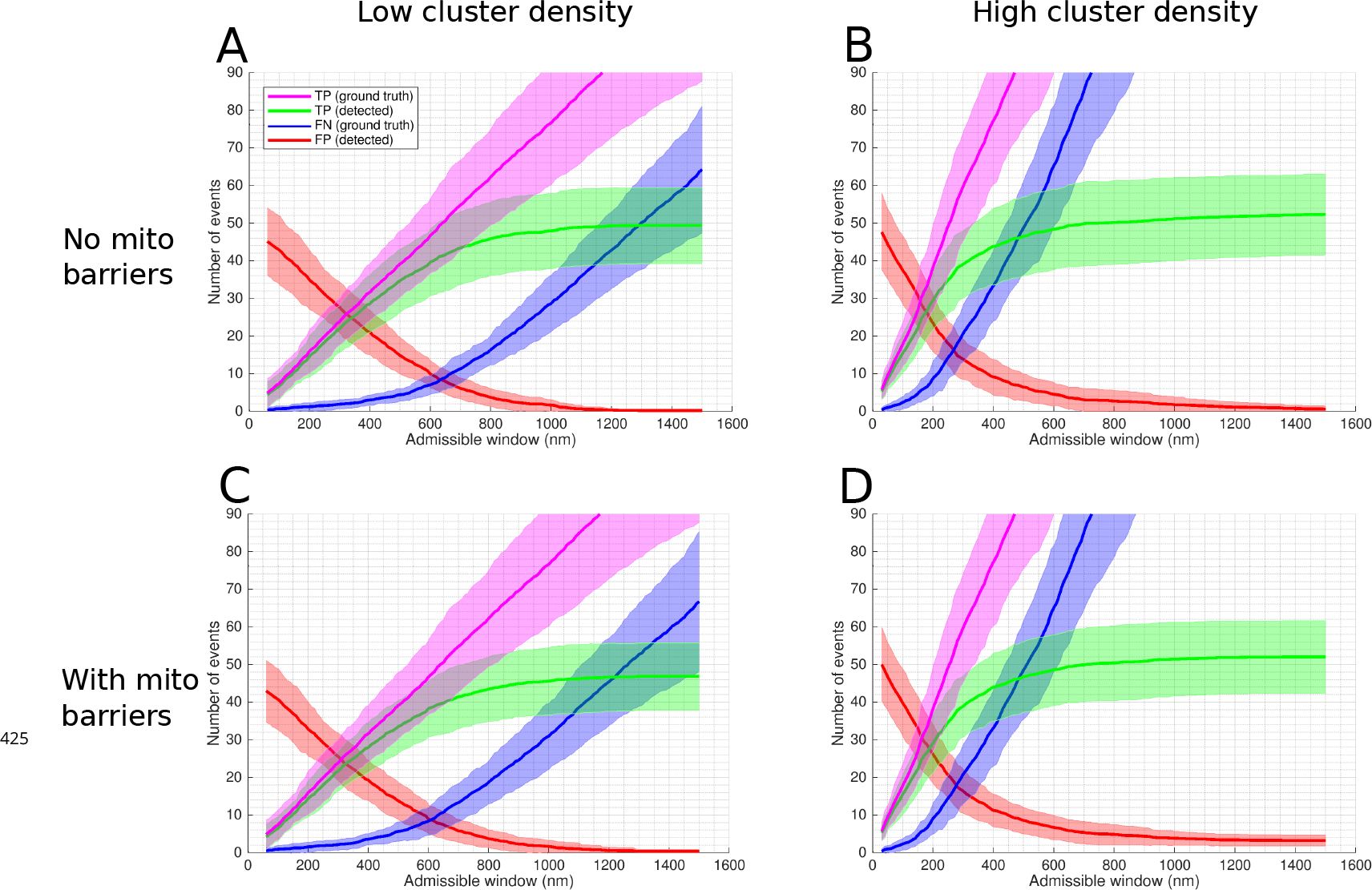
Statistical classifiers of CaCLEAN performance on simulated microscopy data in four model permutations. Statistical classifiers for CaCLEAN results shown as a function of the size of the admissible window for each of the four model permutations. In each case, 22 images were simulated with equidistant spacing through the model volume. Shaded regions indicate one standard deviation of values within the group of 22 image slices. (A) A minimum spacing of 1 *μ*m is enforced between modeled clusters, no mitochondria present in the volume (resulting in a continuous domain across the mitochondrial regions). (B) Cluster spacing is determined by statistical analysis of RyR cluster distributions from immuno-labelled super-resolution microscopy data, no mitochondria present in the volume. (C) A minimum cluster spacing of 1 *μ*m is enforced between modeled clusters, mitochondrial regions are subtracted from the volume, acting as barriers to diffusion. (D) Cluster spacing is determined by statistical analysis of RyR cluster distributions from immuno-labelled super-resolution microscopy data, mitochondrial regions are subtracted from the volume, acting as barriers to diffusion.

**Figure 5–Figure supplement 1.**
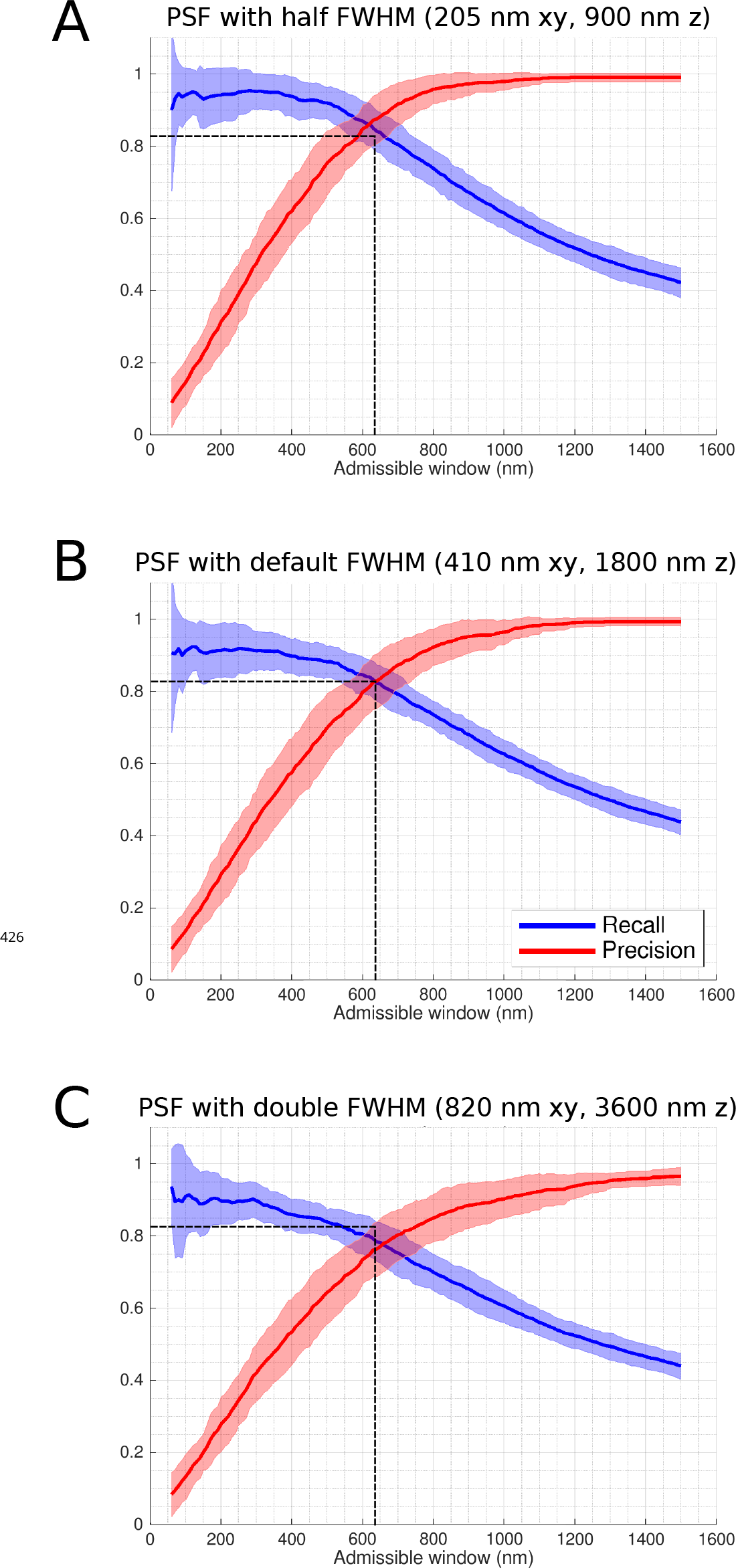
Recall and precision with alternative PSFs. Recall and precision shown for the low cluster density, 1 *μ*m minimum spacing case (Figure 5A), with two alternate point spread functions (PSFs) shown in comparison to the default. The default PSF full width at half maximum (FWHM) dimensions of 410 nm in the image x-y plane and 1800 nm in the through-image z plane were based on published values using a Visitech confocal microscope (Plumb et al., 2015). (A) Recall and precision with FWHM dimensions halved. (B) Results with the default settings. (C) Results with FWHM dimensions doubled. The default setting intersection point indicator dashed lines are left for visual reference.

**Figure 7–Figure supplement 1.**
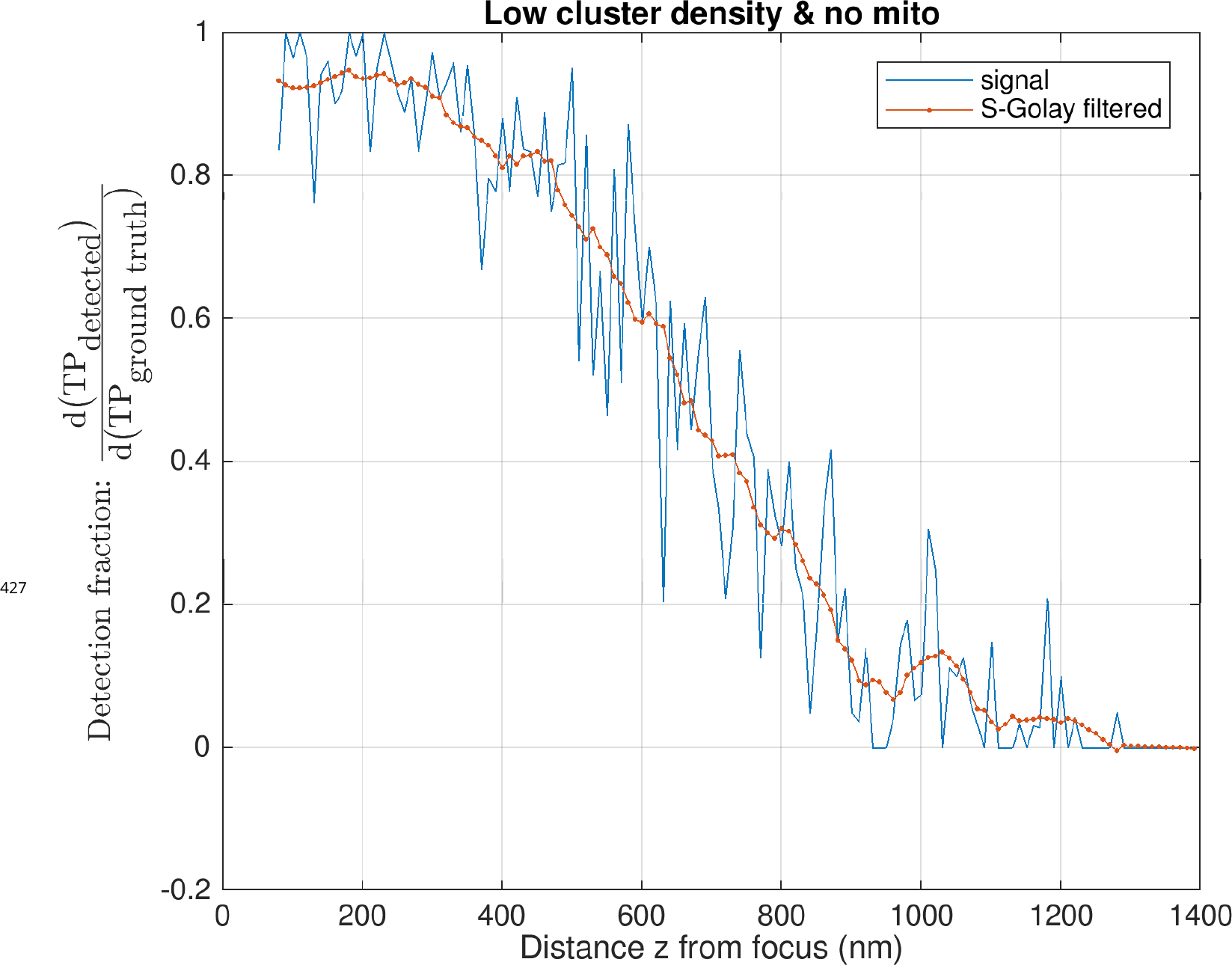
Smoothing filter applied to detection fraction data. The true positive detection fraction results (shown here in blue) were smoothed using a Savitzky-Golay filter (polynomial order 3, frame length 21, results shown in red). The example data shown here is from the low density, no mitochondria case, with the filtered results here corresponding with the magenta data points in Figure 7.

## Footnotes

1 Where the two-dimensional simulated image plane represented x-y axes.

2 True negatives were not considered in this application, as they would represent the set of all remaining locations that were neither modeled nor detected.

3 Immuno-labelled confocal data populations may still slightly underestimate cluster density owing to diffraction-limited resolution.

4 See Figure 3Cb in Tian et al. (2017)

